# Coalescent models for developmental biology and the spatio-temporal dynamics of growing tissues

**DOI:** 10.1101/022251

**Authors:** Patrick Smadbeck, Michael PH Stumpf

**Affiliations:** Centre for Integrative Systems Biology at Imperial College London, London, United Kingdom

## Abstract

Development is a process that needs to tightly coordinated in both space and time. Cell tracking and lineage tracing have become important experimental techniques in developmental biology and allow us to map the fate of cells and their progeny in both space and time. A generic feature of developing (as well as homeostatic) tissues that these analyses have revealed is that relatively few cells give rise to the bulk of the cells in a tissue; the lineages of most cells come to an end fairly quickly. This has spurned the interest also of computational and theoretical biologists/physicists who have developed a range of modelling – perhaps most notably are the agent-based modelling (ABM) — approaches. These can become computationally prohibitively expensive but seem to capture some of the features observed in experiments. Here we develop a complementary perspective that allows us to understand the dynamics leading to the formation of a tissue (or colony of cells). Borrowing from the rich population genetics literature we develop genealogical models of tissue development that trace the ancestry of cells in a tissue back to their most recent common ancestors. We apply this approach to tissues that grow under confined conditions — as would, for example, be appropriate for the neural crest — and unbounded growth — illustrative of the behaviour of 2D tumours or bacterial colonies. The classical coalescent model from population genetics is readily adapted to capture tissue genealogies for different models of tissue growth and development. We show that simple but universal scaling relationships allow us to establish relationships between the coalescent and different fractal growth models that have been extensively studied in many different contexts, including developmental biology. Using our genealogical perspective we are able to study the statistical properties of the processes that give rise to tissues of cells, without the need for large-scale simulations.

## Introduction

The connection between space and time is fundamental to developmental biology. For over a century the location of stem cell proliferation and differentiation during development has been known to be well-organized and of paramount importance to cell fate decision making (e.g. Spemann organizer and primitive knots) [1]. Through the control of cell division and other cellular action, spatiotemporal chemical signalling form complex patterning vital to proper tissue development [2, 3]. Despite the long-established importance of spatial information in understanding tissue development, it was not until relatively recently that widespread understanding of these effects has been possible.

More recent experimental work (relying on advanced microscopy [4] with suitable dyes [5] and fluorescence tags [6] etc.) in the context of developmental biology has focused on cell tracking and lineage tracing. These experiments have already given rise to profound new insights. While certain organisms (such as, most famously, *C. elegans*) are essentially deterministic in their spatial organization because of highly regulated developmental processes [7], most organisms are not so predicable. Opacity, three-dimensional effects, and stochasticity all make lineage tracing and cell tracking experiments difficult [5, 8]. Even if supported by state-of-the-art computational and statistical analyses, these experiments will remain challenging. Computational modelling is therefore emerging as a desirable, and ultimately essential tool to understand the carefully orchestrated processes underlying tissue growth and homeostasis. Modelling is encroaching on biological territory that had previously been predominantly qualitative and descriptive because: (i) sophisticated computational and statistical approaches are required to extract, handle and interpret the incoming data; and (ii) mathematical or computational models can encapsulate complicated and quantitative mechanistic hypotheses and be used to test systematically which aspects of these hypotheses are borne out by reality.

Modelling of tissues has a long-standing history; while reaction diffusion approaches continue to be useful and still dominate much of the literature, agent-based models (ABM) are gaining in popularity. ABMs allow us to include the cellular composition of tissues from the outset. Like cells, agents interact with their environment and each other; occupy finite spatial areas/volumes; and can exhibit the hallmarks of cell behaviour: differentiation, proliferation, movement, and death. All of these factors lead to cells organizing themselves into tissues. While there is a large deterministic component underlying tissue growth (as well as homeostasis), experiments tracing cells and their progeny often demonstrate substantial variability in the lineage behaviours, easily captured by ABMs. Thus, ABM approaches provide a natural computational complement to lineage tracing experiments making it possible to make comparisons to analogous computational models of tissue growth, development or stem cell dynamics.

As cells produce offspring differences can emerge. Some cells produce more progeny than others with many cells producing no off-spring at all. This competition for space and resources will eventually result in progeny from one or very few ancestral cells dominating the tissue (or section of tissue). Cheeseman et al. [16] have denoted dominant ancestral cells as *superstars*; these authors developed ABMs that describe the neural crest growth and development and found that stochastic effects and competition between cells for positions in the new layers of growing tissues appear to affect the development of the enteric nervous system. Similar phenomena are reported from other lineage tracing studies in both healthy and malignant tissue growth [17–19].

There are, however, many unaddressed or poorly understood issues as to how ABM approaches are best related to lineage tracing data. The most glaring issue is that of computational load. To physically observe a biological system it must be comprised of billions of cells. Simulating a single example of such a system may take hundreds of CPU hours to complete. Stochasticity and applications such as Bayesian inference that require hundreds or thousands of such simulations can be all but impossible to implement without incurring massive computational cost. Fortunately, by implementing methodologies inspired by population genetics it may be possible to distil many important lineage tracing concepts into easily digestible and computationally light quantitative rules instead.

When we trace the ancestral relationships among the cells in a tissue we recover genealogical relationships that are familiar from population genetics. This similarity — which is more than just superficial — will be exploited here. In population genetics we study stochastic processes that describe the ancestral relationships among alleles in a population. Population genetics has been applied with great success to, for example, map out the genetic history of human and animal populations, estimate the age of alleles (as well as likely geographical origins), and map out past population movements. A sophisticated mathematical framework has been developed that allows us to elucidate evolutionary dynamics; for example in a population of *N* alleles that evolve according to the standard neutral model, the time until an allele becomes fixed is 2*N* generations (with a variance of *N* generations). One of the reasons for the success of population genetic theory is that evolutionary processes typically occur over time-scales that are so long that they can (with the exception of long-term evolution experiments in bacteria) not be observed experimentally. Instead mathematical models are used to capture the evolutionary dynamics and relate them to the observed data using statistical methods — it is probably no coincidence that evolutionary theory has been linked in lock-step to developments in statistical theory and practice.

One of the fundamental insights that has made this connection between evolutionary/population genetic theory and statistics even tighter is the realization that we can reconstruct the genealogical processes underlying a sample of alleles (drawn from a large population); i.e. we do not have to model the evolution of a large population of *N* individuals forward in time, but can instead look at the stochastic process that describes the ancestral relationship among *n ≤N* (typically *n ≪N*) individuals/alleles [13]. Starting from the present sample we follow their ancestral lineages back in time until all lineages have coalesced into a single lineage; this allele/state is called the most recent common ancestor (MRCA). In addition to the computational efficiency (compared to forward simulations) this *coalescent approach* also focuses explicitly on the observed data and the properties of the underlying genealogical process.

Here we will adapt and apply coalescent theory to developmental processes. Coalescent theory allows us to study populations of cells and their ancestral relationships backwards in time and space. In fact it is the relationship between space and time that comes to the fore in this framework. Developmental processes organize cells into tissues by controlling their cell fate decision making in both space and time. The perspective developed here reveals how space and time are related for different tissue growth models. We will also establish a connection between the neutral evolution model typically explored using coalescence and evolution through fractal growth systems typical of more self-organizing systems. By forging this connection it becomes possible to exchange methods and results obtained through either coalescent theory or the extensive body of work dealing with more complex fractal growth systems.

Here we consider three different models: (1) layer-by-layer growth of a tissue (where we will highlight the relation to a neutrally evolving, constant-sized population [21, 22] and exact coalescent models [23]); (2) asynchronous cell-by-cell growth with diffusion [16] (which is related to the classical Eden growth model [24]); and (3) asynchronous and unbounded growth without diffusion as a bacterial colony or 2D tumour growth model (akin to Sottoriva et al. [20], an Eden growth process). We develop these models in the necessary detail in the *Methods* section.

The *Results and Discussion* sections will then touch on three primary goals. First, to establish that there is a predictable relationship between space and time in both bounded and unbounded (bacterial colony or tumour growth) tissue growth models. Second, to show genealogical or retrospective approaches are applicable to tissue growth and developmental processes. Third, that we can determine scaling relationships that link the very simple coalescent models with more complicated fractal growth models. The ability to consider developmental processes from a coalescent point of view is an attractive complementary approach to lineage tracing. While lineage tracing focuses on the fate of a few chosen lineages — those that will persist for long times as well as those that go extinct swiftly — these are characteristics that inform the coalescent process. More generally, however, our analysis and results show that complicated as well as complex^1^, self-organizing systems can be understood in relatively simple terms once we condition on the outcome, in this case the existence of a tissue with a specific underlying growth model. As we will argue in more detail below a rescaling of the dynamic scale (time and/or space) often suffices to draw out the similarities.

## Methods

The two general models covered in this study are the exact coalescent (related to the Wright-Fisher model [23] of population genetics) and tissue growth models that are inspired by or related to the classical Eden model [24] (or more general models such as the processes described by the KPZ equation). This background section is split into two parts: (1) a basic description of the Wright-Fisher model and exact coalescence along with some of the relevant detailed statistical properties; (2) an algorithmic description of the Eden growth model, the Diffusive-Eden growth model used by Cheeseman et al. [16], and the altered Eden model used to describe tumour growth.

### Coalescent Process

The coalescent is an efficient description of how population samples evolve under a Wright-Fisher model [21] (WFM), which underlies much of population genetics. The WFM is arguably the simplest description of evolutionary change in a population of identical individuals. At each time step (time is measured in non-overlapping generations) the entirety of the population (size *N*) is replaced by choosing random members of the current population to reproduce to form the next generation. The obvious advantage of such a model is its straightforward probabilistic description whereby each member of the population is equally likely to be the parent of any child in the next generation (neutral evolution).

Population genetics, however, generally seeks to obtain information about a population’s history based solely on a current population’s genetic data. Coalescent theory attempts to reverse time and explore a population’s history by tracking how distinct lineages (branches on a family tree) eventually combine (coalesce) as the population is traced backwards in time. With this in mind Kingman, in 1982 [22] (Griffiths [25] and Tajima [26] published their near-identical approaches almost simultaneously), was able to use the Wright-Fisher model and reverse time to achieve exactly this via a genealogical or coalescent approach. Modern population genetics, including the interpretation of, for example, the HapMap data and the determination of a most recent common ancestor (MRCA) of humans, is based on coalescent theory and its derivatives.

With coalescent theory time is reversed — the present is *t* = 0 — and the number of relevant lineages (*n*) that are ancestral to the present-day sample are tracked backwards through time until the MRCA has been reached. These lineages can coalesce (when two lineages arrive at their most-recent common ancestor); this death-process in the number of active lineages can be modelled probabilistically as can the times spent with a specific number (*k*) of active lineages (*T*_*k*_). The key to the use of coalescent theory in population genetic analysis is in its limiting behaviour. At several common limits (large population, *N→ ∞*, and small relevant initial lineage number, *n ≪ N*) the coalescent process is characterised by an exponentially distributed reaction process, where the time before the next coalescent event among *k* lineages, *T*_*k*_, can be determined using a uniform random number, *u ∼ U* (1), as

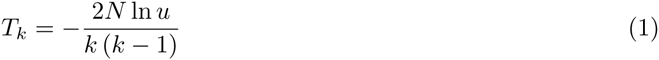

Here *T*_*k*_ is the time spent with *k* lineages. Overall, starting at a full population (*n* = *N*) the total time to the most recent common ancestor (*T*_MRCA_) can be approximated as:

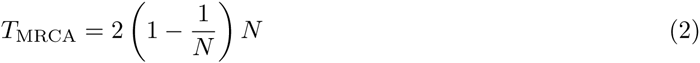

A full derivation and explanation of the results provided here is found in any book concerning coalescent theory and population genetics [27]. While the Kingman coalescent uses this large population approximation (and in particular *n≪ N*) an exact description of the Wright-Fisher model is provided by Fu [23]. The fundamental difference between the two is that in conventional coalescent not more than one coalescent events can only occur in any generation. The exact coalescent or Λ-coalescent provides an exact description of the genealogy of a population (rather than a small sample) where multiple coalescent events can occur in a single time step. In this study we exclusively use the Wright-Fisher model, which we call the neutral evolution model. When the starting population is large (here *N >* 100) both Equations 1 and 2 are applicable.

Coalescent theory has been developed into many different directions and can account for genealogical processes in populations of variable sizes. The wealth of knowledge amassed in the coalescent theory literature allows us to apply our analysis also to the bacterial colony/tumour growth model; in this case the statistics provided in Equations 1 and 2 do not apply, although results for such models can be found in the literature [27–31].

### Eden Growth Models

In Eden growth models, instead of evolving populations of individuals, we are looking at tissues growing over time — it is for these processes that we seek genealogical representations of the ancestry of cells. We are motivated by a tissue growth model introduced by Cheeseman et al. [16], probably more precisely described as an Eden growth process [24] with diffusion. The Eden model has been a stalwart of analyses into surface and bacterial colony growth and forms a generic example of a fractal surface on a lattice first described in 1961 [24]. The basic assumptions are that the growth occurs at the boundary of the growing system, and that the system remains connected. While variations have arisen (often to facilitate computational analysis [32]), the basic algorithmic growth process can be described by Algorithm 1.

#### Algorithm 1 Eden model

~~~
Initialize set of *N*_0_ cells *A*, *N* = *N*_0_ on the lattice
Determine set of boundary sites, *B*, from *A*
**while** *N ≤N_max_* **do**
   choose site *b* from boundary sites *B*
   add cell *b* to set *A*
   recalculate set of accessible boundary sites *B* from *A*
   *N*= *N* + 1
**end while**
~~~

Eden growth is a good, generic yet qualitatively accurate description of many growing systems in nature (including cellular systems) and it has been used extensively across many different fields [33–35]. The analysis of fractal growth systems from a coalescent perspective could bridge the gap between the simplicity of the coalescent and the complexity of realistic biological systems. Unlike in neutral evolution for population growth models the cells in fractal growth systems compete for space and thus incorporate a form of selection. Selection and asymmetric growth (cellular generations overlap) are two fundamental differences between neutral evolution and fractal growth models that complicate the use of coalescent theory in spatial lineage tracing studies.

In Figure 1A we provide a graphical representation of a single step during Eden growth. Figure 1B shows the same single step for the coalescent (Wright-Fisher) model for comparison. In Figure 1C an example of lineage tracing (and coalescence) in the Eden model (N = 20) is shown starting at the 200th generation and tracked from parent to child backwards in time. The most recent common ancestor (MRCA) is marked in black, the generation in which there exists only a single surviving lineage. The time to the most recent common ancestor (*T*_*MRCA*_) is thus measured in generations backwards in time.

In several publications by Landman and colleagues [16, 36], a slight modification of the Eden growth model was introduced. The model was intended to describe the growth of the neural crest during embryonic development and was originally presented on a cylindrical surface with growth in the z-axis and periodic boundary conditions. The model by Cheeseman et al. [16] differs from Eden growth by incorperating diffusion whereby cells are able to move in addition to reproduce. This model is described in Algorithm 2. The non-diffusive version of this model is a true Eden growth model and was used in bacterial colony growth simulations, shown in Algorithm 3. Note that in these cases the simulation was stopped after a specified generation threshold had been reached.

In total three distinct models are used to simulate tissue growth in this study, and we will relate their behaviour to principles from coalescent theory. The Wright-Fisher model is used with an exact coalescence for comparison. The biological growth models are then simulated with a more accurate Diffusive-Eden model for unidirectional tissue growth, and a true Eden growth model for bacterial colony (or two-dimensional tumour) growth.

**Figure 1.**
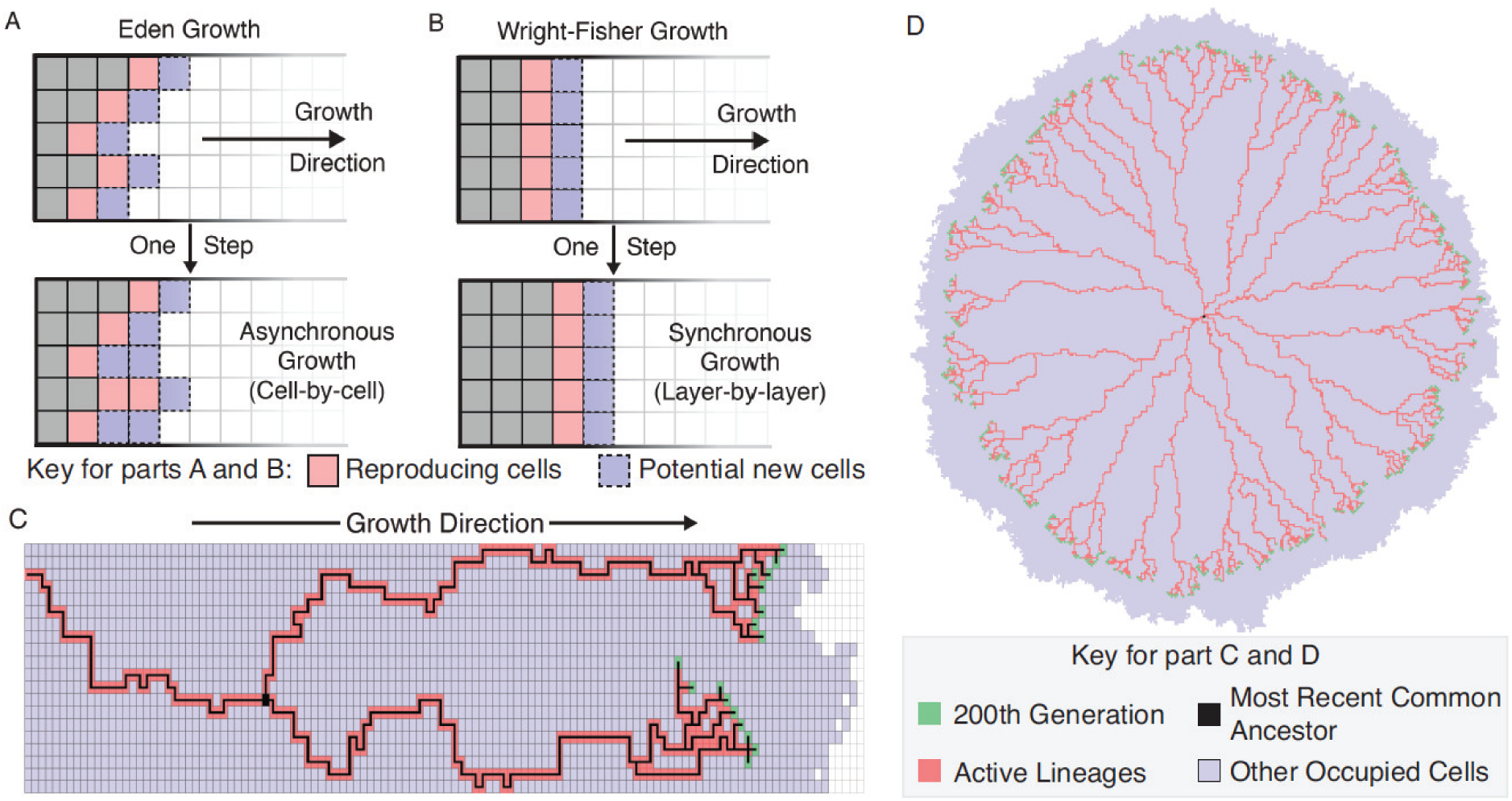
Example of Wright-Fisher and Eden results for different geometries. **(A) A** graphical description of a single step in a small Eden model. Note that the potential new cells (blue) are chosen proportional to the number of reproducing cells (red) that surround the site. Parents are also chosen at random from the neighboring reproducing sites. This is an asynchronous process as new layers in the tissue growth model can begin before the previous layer is complete. (B) A graphical description of the Wright-Fisher growth process. Here the reproducing cells are chosen at random to be the parent of a cell in the next generation. This growth process is synchronous as all cells in the next generation and next layer are produced at the same time. (C) An example of an Eden growth process on a bounded domain growing in a single (left-to-right) direction. The 200th generation is marked in green, active lineages in red, and the MRCA in black. The MRCA is 104 generations in the past (contain within the 96th generation). (D) A tumour growth model (unbounded, with an initial population of one located at the origin). The 200th generation is marked in green, active lineages in red, and here the MRCA is the initial generation shown in black. This is characteristic of unbounded tissue or bacterial colony growth that exhibit star-like lineages.

## Results and Discussion

The intention of our analysis is fourfold: (1) to forge a direct connection between the space and time dimensions in Eden-like tissue growth models; (2) to establish that the idea of “superstars” as presented by Cheeseman et al. [16] is, in fact, a logical, inevitable, and indeed expected outcome that is readily captured by the coalescent process; (3) to determine the scaling factors for coalescent models of biological growth processes and to establish the link to the classical coalescent (as applied to the Wright-Fisher model); (4) to show that we can establish simple scaling relationships for both unidirectional and unbounded growth systems.

### **Algorithm 2** Diffusive-Eden Model

~~~
Initialize set of *N*_0_ cells *A*, *g* = 1
**while** *g ≤g_max_* **do**
  choose cell *a* from current cell population *A* to diffuse
  choose an adjacent site (*d*) for diffusion
  **if** *d* is unoccupied **then**
     move cell *a* to site *d*
  **end if**
  choose cell *a* from current cell population *A* to proliferate
  *g*_*a*_ is the generation number for the cell *a*
  choose an adjacent site (*p*) for proliferation
  **if** *p* is unoccupied **then**
    generate new cell at site *p*, add to set *A*
    the generation number for the cell *p* is *g*_*p*_ = *g*_*a*_ + 1
    **if** *g*_*p*_ > *g* **then**
        *g*= *g*_*p*_
    **end if**
  **end if**
**end while**
~~~

### **Algorithm 3** Eden Model for Colony Growth

~~~
Initialize set of *N*_0_ cells *A*, *g* = 1
**while** *g≤ g_max_* **do**
   choose cell *a* from current cell population *A* to proliferate
   *g*_*a*_ is the generation number for the cell *a*
   choose an adjacent site (*p*) for proliferation
   **if** *p* is unoccupied **then**
      generate new cell at site *p*, add to set *A*
      the generation number for the cell *p* is *g*_*p*_ = *g*_*a*_ + 1
      **if** *g*_*p*_ > *g* **then**
         *g* = *g*_*p*_
      **end if**
   **end if**
**end while**
~~~

### The Neutral Evolution Model Results

We begin with a “neutral evolution model” whereby tissue growth proceeds in a layer-by-layer fashion with parent cells chosen at random from among the *N* cells in a population. Such a system could be thought of as a simplified version of unidirectional and bounded (constant number of cells in each tissue layer) tissue growth. In Figure 2 we show the coalescent results for three neutral evolution models with tissue widths of *N* = 10 (dots), 100 (solid), 1000 (dashes). The mean number of lineages (red) demonstrate an initial fast decay as a substantial portion of lineages are eliminated in the first few time steps. This bahaviour is characteristic of the exact coalescent where many multi-lineage coalescent events occur per generation. The probability of coalescence (blue) shows the inevitable coalescent process in which nearly all simulations will coalesce within 5 *× N* generations. By scaling the time axis with the tissue width *N*, it is easy to see how as *N* tends to infinity the coalescent process converges onto a single general trajectory characterized by the statistical properties from Equations 1 and 2.

**Figure 2.**
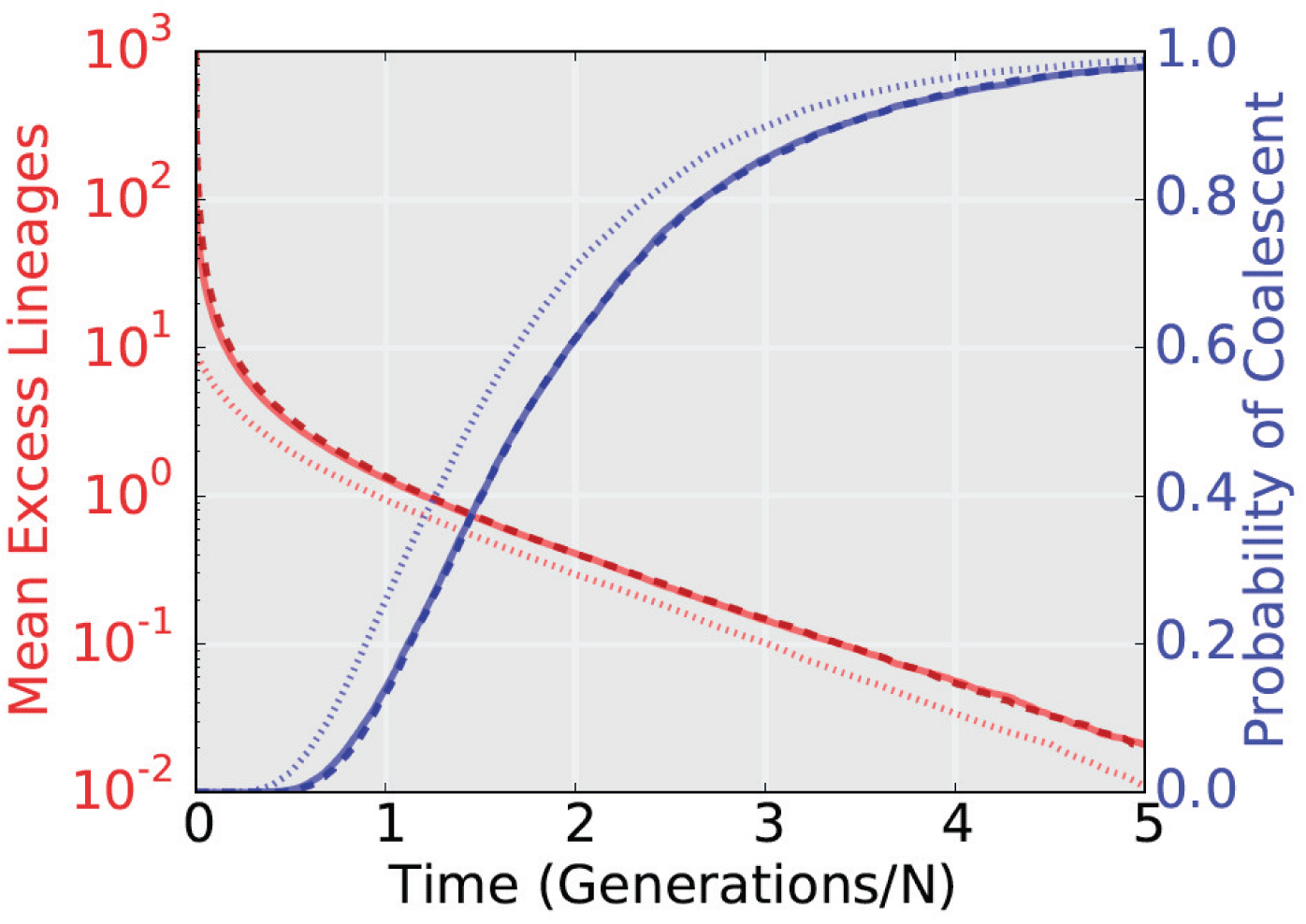
Coalescent properties for layer-by-layer tissue growth. A bounded unidirectional layer-by-layer growth process is shown for three different tissue widths (with *N* = 10 (dots), *N* = 100 (solid), and *N* = 1000 (dashed) cells). The mean excess lineages (defined as the number of lineages in addition to the single commen ancestor of the population) is shown in red on a semilog scale. The probability of having achieved coalescences by a specific cell depth/generation is shown in blue. The time axis is scaled with tissue width (*N*) revealing an asymptotic relationship as *N→ ∞*. All simulation are the result of 10,000 monte carlo simulations.

By using neutral evolution as a stand-in for tissue growth the connection between tissue depth (spatial) and the generation number (temporal) is deterministic. With each added layer the generation number grows by one as the population’s children completely supplant the parent generation. Thus, an analysis based on an exact coalescence (temporal) is directly equivalent to describing cell lineage tracing in a spatial regime.

### The Diffusive-Eden model results

The analysis presented in the previous section becomes less straightforward when using the more realistic Eden growth model. Specifically, the connection between space and time is not immediately apparent as it is in the Wright-Fisher model. In Figure 3A the Diffusive-Eden growth process is shown whereby a distinct spatiotemporal connection is apparent. Generations 1 (red), 50 (yellow), 100 (blue), 150 (green) and 200 (magenta) are shown with each lattice location weighted according to the probability a cell was present in one million stochastic realizations of the Diffusive-Eden growth model. While the distribution along the tissue width quickly reaches uniformity, Figure 3B shows how the distribution along the tissue depth appears normally distributed with an increasing mean and variance.

It should be noted that throughout this study time is represented as a series of successive generations in order to more easily translate between the neutral evolution and Eden growth models. Assuming cell division time scales with the number of available boundary sites (e.g. each cell divides independently by an exponential distribution) this is appropriate.

**Figure 3.**
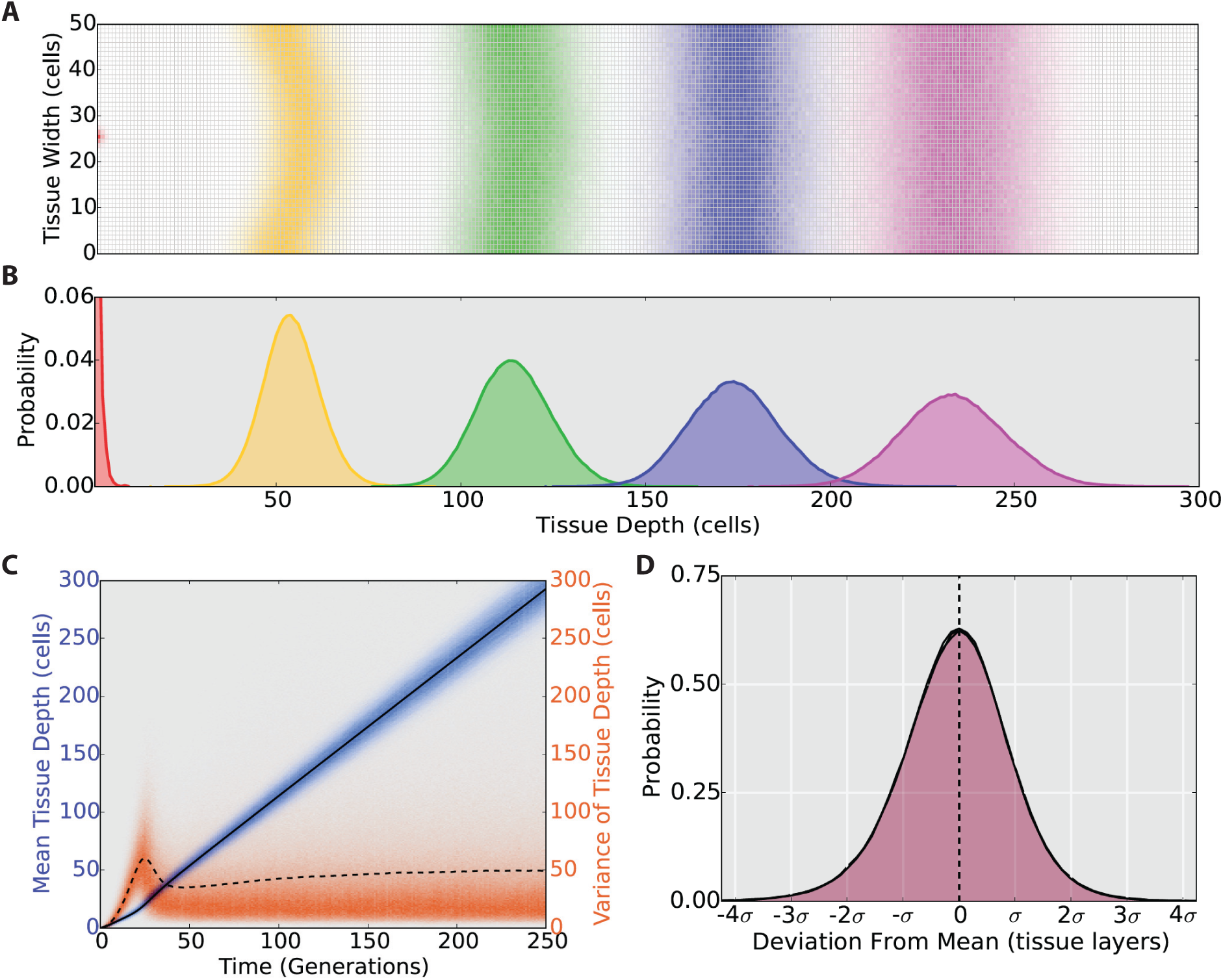
Spatiotemporal connections in the Diffusive-Eden model. (A) Probabilistic representation of the position for generation 0 (red), 50 (yellow), 100 (green), 150 (blue) and 200 (magenta) in 1 10^6^ Monte Carlo realizations of the Diffusive-Eden growth model. Each lattice point is weighted according to the probability that a cell of the specified generation would occupy that site during simulation. (B) A one-dimensional representation of the probability distributions for site occupancy across the tissue depth. (C) Mean tissue depth (blue distribution) and the variance for the tissue depth (orange distribution) versus time as represented by the cellular generation number. The overall mean (solid line) and mean variance (dashed line) for 1 10^6^ simulations is also shown. Note that the variance is a long-tail distribution and thus the mean is significanly higher than expected given the distribution. (D) The deviation from the mean tissue depth for generations 250 (red), 200 (blue, obscured), and 150 (green, obscured) calculated from 1 · 10^6^ simula*v*ti<u>on</u>s. Note the distribution appears normally distributed with variance equal to *N* (tissue width, 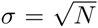) and stationary.

The increasing variance, however, is in actuality a function of the mean tissue depth for a particular cellular generation as shown in Figure 3C. Here the mean tissue depth versus generation number (blue distribution, solid black line shows the mean) shows a normally distributed function with an increasing mean and variance. Alternatively, the variance around this mean (orange long-tailed distribution, dashed black line showing the mean value) quickly reaches a steady state. The values were calculated by taking the *N*_*g*_ cells in generation *g* in any particular simulation and obtaining a mean and variance for the tissue depth for the group. This suggests that there is an easily defined statistical relationship between time (generation number) and space (tissue depth) and thus, if coalescence in time can be determined, the statistics for spatial cell lineage tracing would be readily determined.

Indeed, as shown in Figure 3D, there is a steady state distribution for the deviation from the mean position for any cell (of a specific generation) during an individual stochastic realization of the Diffusive-Eden model. The three overlapping distribution are normally distributed around zero with a variance equal to *N* and very quickly reach a stationary distribution. Using two normally distributed random numbers a statistically accurate location for a cell of generation *g* can be determined for this Diffusive-Eden growth. The values were calculated by taking the *N*_*g*_ cells in generation *g* in any particular simulation and obtaining the deviations from the mean tissue depth for the group. This relationship is related to the underlying fractal growth dynamics for the Eden model whereby the roughness of a unidirectional Eden growth frontier is proportional to *N* ^1/2^ [37].

The connection between space and time in a realistic mathematical treatment of a biological growth process suggests an opportunity to expand the influence of coalescent theory beyond population genetics and into cell lineage tracing in developmental biology. By knowing how far back in generation number coalescence is likely to have occurred one can approximate a statistically valid spatial location for the MRCA in a growing tissue using a relationship much like in Figure 3.

The next question concerned whether the “superstars” observed by Cheeseman et al. [16] were a manifestation of inevitable coalescent as in the simpler neutral evolution model. In Figure 4A the lineage numbers (red) and probability of coalescent (blue) are shown for tissue widths of *N* = 10 (dotted line), 50 (solid line), and 100 (dashed line).

**Figure 4.**
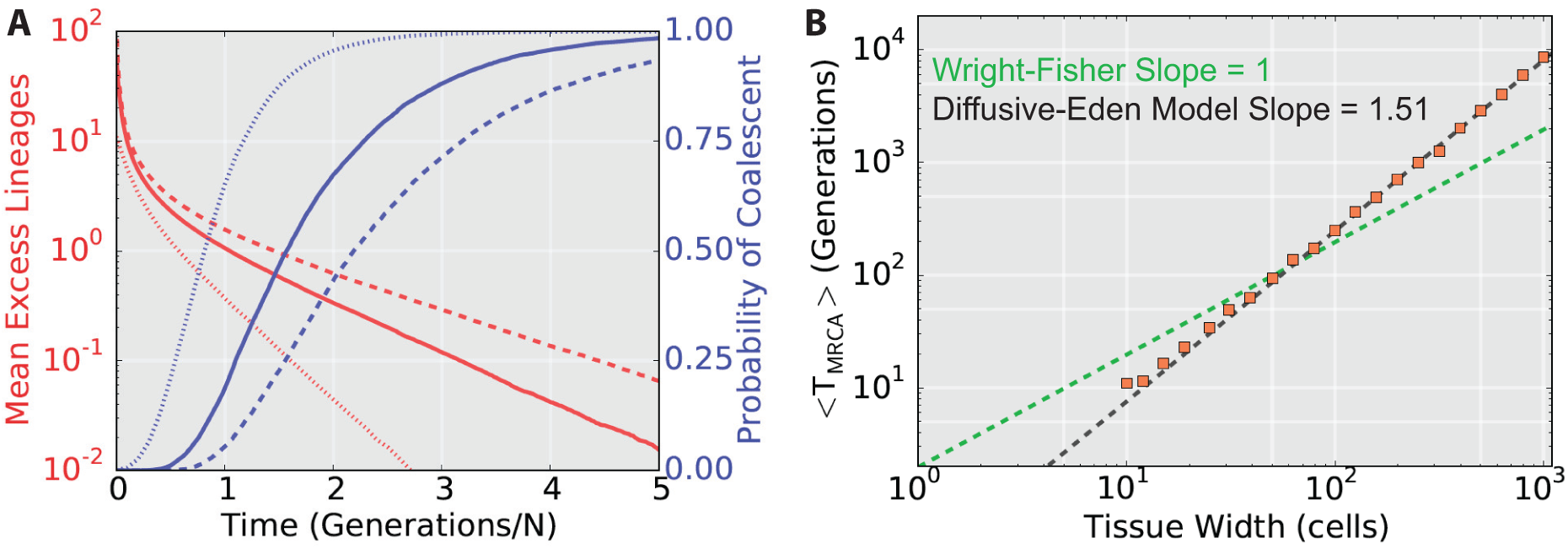
Coalescent analysis of the Diffusive-Eden model with time to the most recent common ancestor scaling. (A) The mean excess lineages (defined as the number of lineages in addition to the single commen ancestor of the population, red) and the probability of coalescence (blue). Results are shown for three tissue widths for a Diffusive-Eden model, *N* = 10 (dots), 50 (solid line), and 100 (dashed line) from 10,000 Monte Carlo simulations. (B) The *T*_*MRCA*_ for the Diffusive-Eden model is shown for tissue widths from *N* = 3 to 1000 (red squares, 1000 instances simulated) on a log-log plot. A comparison line for the Kingman coalescent (based on Equation 2) is shown as a green dashed line for reference. A linear regression to the Diffusive-Eden results is shown as a dashed black line and exhibits a calculated slope of approximately 1.51. These linear results are used as a basis for the effective population in Equation 3

These results show two important characteristics for the Diffusive-Eden model: first, the coalescence for the results from Cheeseman et al. [16] were inevitable (in that study *N* = 50) for the tissue depth being simulated; more importantly, however, in the limit (*N→ ∞*) this will not be the case. As *N* increases the time to coalescence will drift further into the past until coalescence will not be found under reasonable time/tissue depth expectations. This suggests that while the superstar model may be important on developmental spatial scales, the process itself does not necessarily apply to larger systems. Instead large tissues must be founded by several stem-cells which will in turn likely derive from more general stem cells upstream in the differentiation hierarchy.

The results from Figure 4A beg the question as to what the scaling properties (with respect to the size of the tissue) of the *T*_*MRCA*_ are for the Diffusive-Eden model. In Figure 4B a log-log plot for the average *T*_*MRCA*_ for tissue widths from *N* = 3 to 1000 is shown (red squares, 1000 simulations). The time to the most recent common ancestor (*T*_*MRCA*_, Equation 2) for the Kingman coalescent scales with *N* (producing the stationary trajectory on the normalized scale in Figure 2), and is provided for comparison (dashed green line). As is apparent the scaling (slope) of these two lines are very different. A linear regression (black dashed line) can be matched to the Diffusive-Eden data to reveal that the slope of the data is approximately 1.51.

The final question then is whether this scaling factor can be used to draw a connection between the neutral evolution results and the Diffusive-Eden results. A connection could potentially be utilized to allow for a time-based coalescent analysis to be applied universally to tissue or developmental simulations. In Figure 5 the scaling property determined from Figure 4B can be applied to the coalescent results to regain, on average, neutral evolution dynamics. Thus Diffusive-Eden growth is related (via a simple scaling relation) to the classical neutral evolution process. With this in mind we can apply coalescent theory to model the ancestral relationships among cells in developing tissues.

In Figure 5 the time domain for the lineage results (blue) and probability of coalescence (red) from Figure 4A are rescaled using the *N* ^3/2^ factor determined in Figure 4B. These results now match the exact coalescent trajectory on average, and the coalescent probability is nearly identical with increasing *N*. Like with many empirical application of coalescent theory [38] this can be thought of as an effective population number. By scaling by the following effective population:

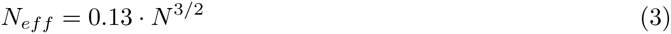

We are able to obtain what is an approximation to neutral evolution coalescence in a more realistic biological growth model for any desired tissue width (*N*).

The different scaling property for *T*_*MRCA*_ for the Diffusive-Eden model is rather striking, but ultimately unsurprising. Primarily because a similar scaling factor plays a prominent role in the underlying dynamics of the Eden model, namely the dynamic exponent for Eden growth *z* = 3/2. Indeed, research in the field of directed polymers suggests that the time to the most recent common ancestor will scale according to *N*^*z*^ (written as *N* ^1/*v*^ in Reference [39]). And it is known that the directed polymer dynamics at zero temperature are equivalent to Eden growth [40]. This result has two immediate consequences for biology. First, by knowing the dynamic exponent (*z*) it suggests that a general scaling rule can be applied *a priori* to a biological system. Dynamic exponents have already been experimentally determined for bacterial colonies, e.g. for bacterial colonies. Second, and more importantly in our opinion, it confirms that the treatment of space as equivalent to (developmental) time is meaningful. Even in a fairly realistic growth model in which cells can move and rearrange, the genetic tree exhibits the same timescale for a spatial domain.

Coalescent theory then is immediately applicable and valuable to the analysis of biological growth models. The emergence of “superstars”, for example, could have been postulated prior to the simulation presented by Cheeseman et al. [16] if the scaling property shown above had been known. Additionally, the presence of superstars for large domain sizes can now be completely ruled out for the Diffusive-Eden model because as *N* goes to infinity “superstars” will eventually never be observed on a practical tissue depth. Going even further the presence of superstars can now potentially be ruled out for a range of tissue models prior to simulation because any fractal growth system with a dynamic exponent larger than one will not observe superstars as *N* goes to infinity. Finally, we can begin to make statements about systems that might otherwise be impractical to simulate. For example, using the dynamic exponent, practical statements might be made about three-dimensional tissue growth models without simulation; for example about the number of stem cells required to generate a tissue of given size in a certain time-frame. These four examples show how coalescent theory may be used to interpret, explain and predict biological results that can be extracted from lineage tracing experiments without the need for large scale simulations (or eventually perhaps even time-consuming experiments).

**Figure 5.**
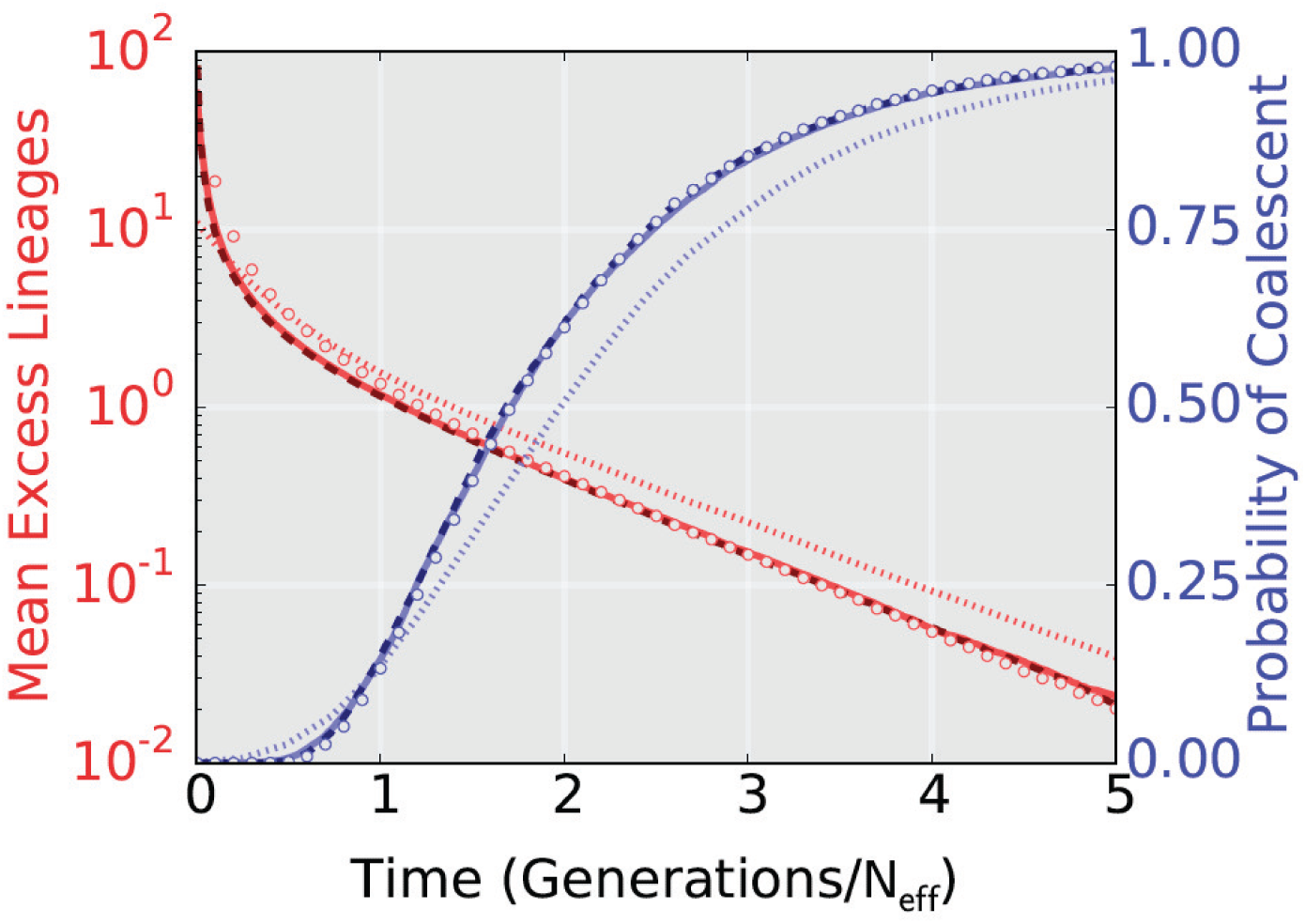
Scaled coalescent analysis for the Diffusive-Eden model results. These results are identical to those presented in Figure 4A, but scaled with an effective population dictated by the results in Figure 4B (*N*_*eff*_ = 0.13 *N* ^3/2^, Equation 3). Again *N* = 10 (dots), 50 (solid line), and 100 (dashed line) are presented. The Wright-Fisher result for *N* = 1000 (empty circles) from Figure 2 is provided for reference for both mean excess lineage (blue) and coalescent probability (red). For the Kingman model specifically *N*_*eff*_ = *N* = 1000 exactly.

### Bacterial colony Eden growth simulation

We next use a true Eden model (non-diffusive, Algorithm 3) without boundary constraints to model a bacterial growth colony that starts from a single occupied lattice point (similar to Figure 1D). The intention is to show that the space-time connection established for bounded growth also holds for unbounded tumour-like growth. Figure 6A shows the occupation probability of different points (weighted according to probability) for generations 1, 50, 100, 150, and 200 as in Figure 3A. Once again the relationship between the generation number (time) and spatial location is readily apparent. Figure 6B shows results grouped according to radial position as well.

In this case the mean position versus generation number (Figure 6C) is noticeably different from that found for the bounded case, Figure 3. The mean radial position is almost deterministic (blue distribution and solid black line), whereas the increasing spread in the generational position data (Figure 6B) can almost entirely be attributed to an increasing deviation from the mean. These results suggest that any individual realization of this stochastic growth process will have a non-stationary distribution as cells in the same generation get farther and farther apart. This increasing spread, however, is predictable and, again, a relationship between time and space can be established for use with a coalescent analysis.

**Figure 6.**
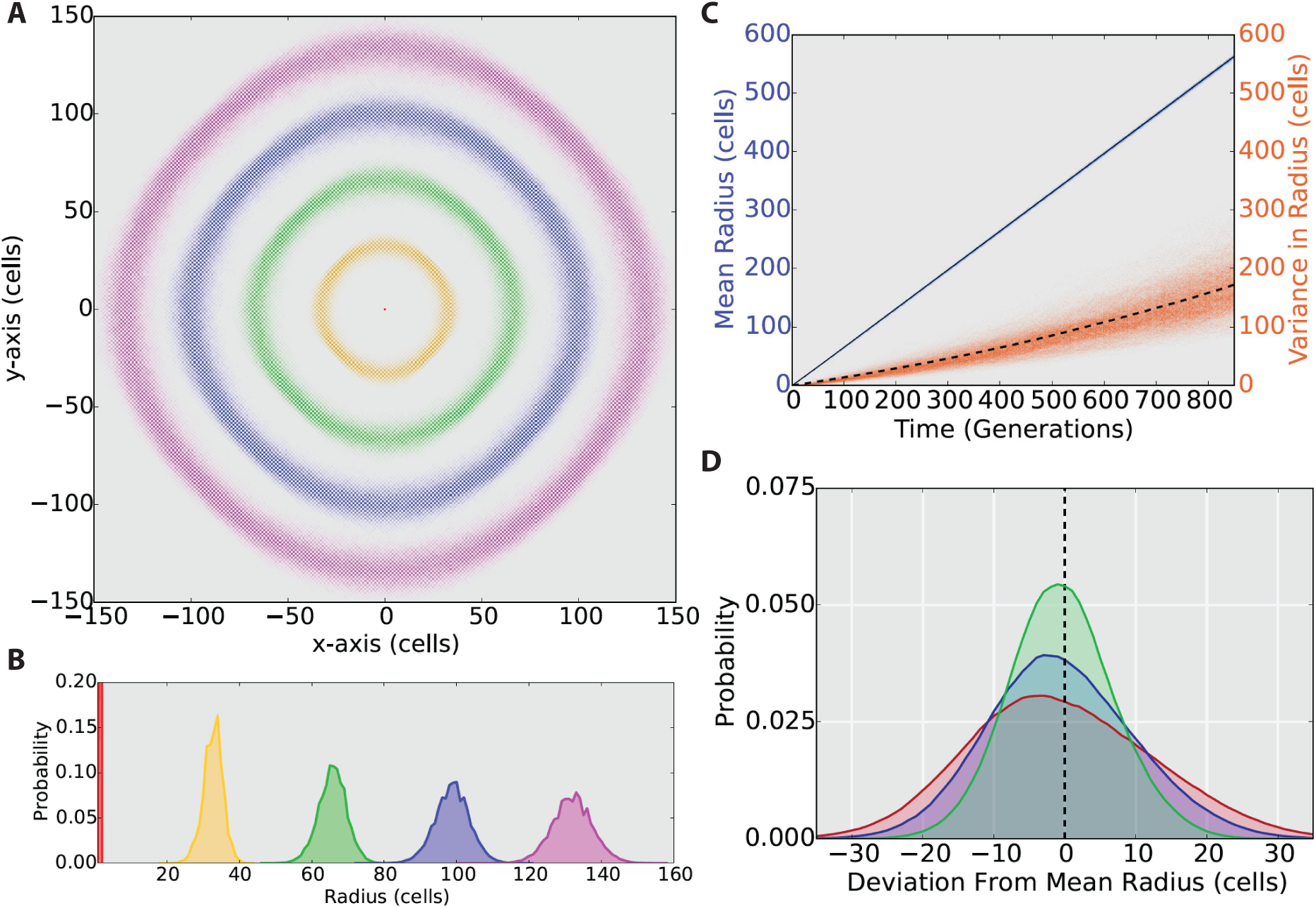
Spatiotemporal connections in a bacterial colony Eden-growth model. (A) Probabilistic representation of the position for generation 0 (red), 50 (yellow), 100 (green), 150 (blue) and 200 (magenta) in 1 10^6^ Monte Carlo realizations of the non-diffusive Eden growth model. Each lattice point is weighted according to the probability that a cell of the specified generation would occupy that site during simulation. (B) A one-dimensional representation of the probability distributions for site occupancy across the radial depth. (C) Mean radial depth (blue distribution) and the variance for the radius (orange distribution) versus time as represented by the cellular generation number. The overall mean (solid line) and mean variance (dashed line) for 1 10^6^ simulations is also shown. (D) The deviation from the mean radius for generations 350 (green), 600 (blue), and 850 (red) calculated from 1 10^6^ simulations.

Indeed, in Figure 6D the distributions for generation 350 (green), 600 (blue) and 850 (red) are shown to diverge as the generation number increases. The lack of a stationary distribution, and thus an ever-increasing roughness to the colony surface, is a well-known consequence of unbounded Eden fractal growth [41]. The lack of a stationary distribution also has an interesting consequence in that the most recent common ancestor is at or close to the originating cell in the system, i.e. *T*_*MRCA*_ is approximately the same as the current number of generations simulated. The number of cells in each generation increases linearly, and with non-constant growth processes a genealogical tree takes on a star-like pattern. The existence of a star-like genealogical tree is well known consequence observed in some tumour growth models [20, 42] and shown in Figure 1D.

A relationship can be found between the coalescent of lineages in a growing Eden cluster and neutral evolution in a growing population through the relationship in Figure 4B. In Figure 7A the number of lineages remaining as the simulation is run backwards in time is shown for generation 450 (solid green line, with *N*_0_ *≈* 1250), generation 900 (solid red line, with *N*_0_ *≈* 2500) and generation 1800 (solid blue line, with *N*_0_*≈*5000) of the Eden growth model. As can be seen a substaintial percentage of the lineages remain when coalescence is forced by the linearly decreasing total population size (gray line). For comparison neutral evolution models run to the same number of generations and an average population growth rate of *V* = 2 · *π* cells per generation are provided for comparison (dashed lines). As can be seen no coalescent occurs in the neutral evolution model as well. While the two lines differ substantially the dynamics of the bacterial growth model and Kingman with linear population growth do appear to have the same basic shape.

**Figure 7.**
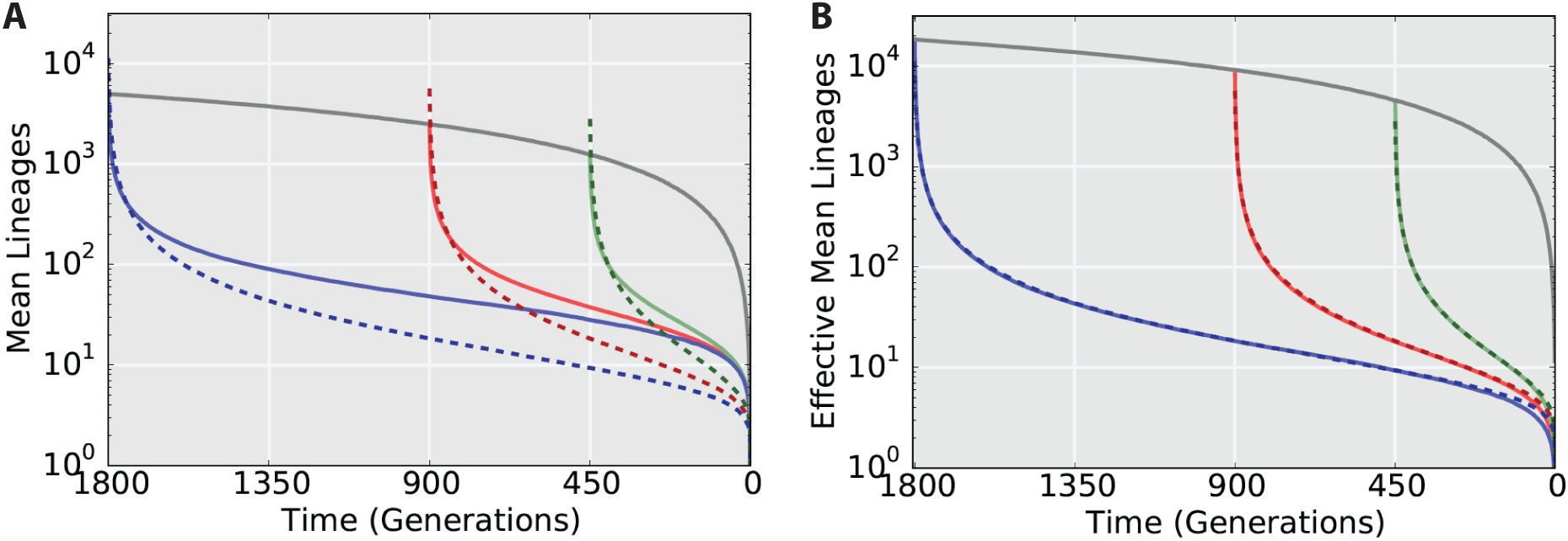
Lineages remaining starting from three generations backwards in time for a bacterial colony or tumour. (A) The Eden growth model for bacterial colony or tumour starting at generation 1800 (blue solid line), 900 (red solid line) and 450 (green solid line) showing the total lineages remaining. The total population (grey solid line) is growing linearly. Coalescence is not observed in almost all simulations (a star-like tree). A neutral evolution model with linear population growth (*V* =2*·π*) is shown as dashed lines for comparison. This model also exhibits star-like trees. (B) Modeling of an effective lineage number remaining now shows strong overlap between the two models. The relationship is presented in the text. All results are calculated from 1000 stochastic simulations.

A relationship between the neutral evolution model and the Eden model results can be established using the same fractal exponents observed in the Diffusive-Eden model results. In 1996 a short paper by Manna and Dhar explored the relationship between the critical exponents of the Eden model and the underlying cell lineage “backbone” (i.e. its genealogical tree if you think of the cluster as a biological system) [43]. The key relationship presented therein was between the fractional number of lineages that survive up to a height *h* away from the original surface, *N*_*h*_:

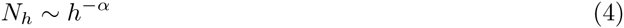

Where *α* = (*d-* 1) */z*, *d* is the dimension of the system (here, we are on a 2D surface), and *z* is the dynamic exponent which for the KPZ system is 3/2 (which we rediscovered via the MRCA in Figure 5). This result can be qualitatively shown to make sense in light of the paper by Brunet, Derrida, and Simon [39], which reports a time of coalescence proportional to *N* ^3/2^ for a two-dimensional system. If we start with *N* cells and coalesce to one cell in *N* ^3/2^ generations, a straight line that connects those two points is proportional to *t*^−2/3^. Since height and time are related linearly (Figure 6C) this exactly corresponds to Equation 4 where *α* = (2- 1) /(3/2) = 2/3.

What we are actually interested in in Figure 7 is the absolute (and not the fractional) number of extant lineages. Thus we can change the equation to,

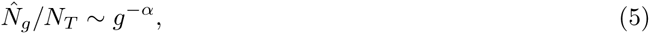

where 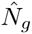 is the absolute number of lineages left *g* generations back in time and *N*_*T*_ is the total lineages at generation *G*-*g* where *G* is the total number of generations considered (or, as here, simulated). *N*_*T*_ is proportional to *G − g* and thus the absolute number of lineages for the Eden system can be written as,

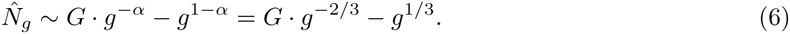

In Figure 7B we relate the Eden results to neutral evolution on a similarly growing population (growing linearly with generation number). Using the above graphical and numerical arguments, this neutral evolution model then scales with an *α* equal to 1 and thus,

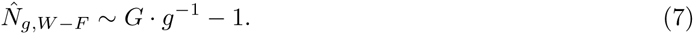

In order to translate the trajectory of the Eden growth model 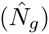 to that of the neutral evolution model 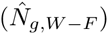 an effective lineage population for the Eden process can be formed,

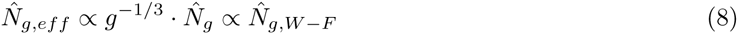

Indeed, in Figure 7B, this simple modification of the Eden growth trajectory results in a nearly perfect overlap with the neutral evolution model with linear growth. The actual modification is as follows:

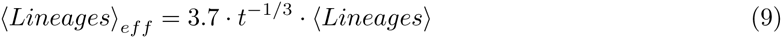

Here time represents the generation number (starting at the first generation in the past). Ultimately the results for the unbounded Eden growth model agree with the results determined through the bounded growth model (and the corresponding coalescent results for growing populations found in Figure 7). These results also confirm the close relationship between the dynamic exponent in fractal growth and the underlying tree structure that is the objective in cell lineage tracing experiments. Both results suggest that it is possible to use results obtained via coalescent theory to capture and describe the results found in complex fractal growth models. In turn statistically valid results, e.g. about the spatial locations of cells belonging to ancestral populations of cells, can be determined from such data and analyses.

## Conclusion

In the preceding sections we have presented three models concerning coalescent theory applications in tissue growth: (1) Layer-by-layer growth using a neutral evolution model; (2) Asymmetric Eden growth with diffusion (akin to Cheeseman et al. [16]); (3) Unbounded Eden growth (without diffusion) as in a bacterial colony or tumour growth model (akin to Sottoriva et al. [20]). Using these models we have established that, in both bounded and unbounded growth, there is a predictable relationship between space and time in realistic tissue growth models. We also showed that coalescence (a dominant “superstar”) is inevitable in Eden growth models on small spatial domains based on coalescent theory, and that there is a non-trivial, but useful, relationship between the Eden growth model and neutral evolution model in unbounded growth. We were able to determine a scaling factor associated with the coalescent processes that are connected to these tissue growth models; this turns out to be scaling factor that is equivalent to the dynamic exponent in fractal growth models (an interesting result that had not been previously shown to our knowledge). This establishes a link between developmental and tissue growth processes on the one hand, and fractal surface growth models on the other. Overall, the results presented herein offer a way to relate spatial cell lineage tracing to the temporal coalescent theory, and similarities are drawn between the well-established and efficient statistical analysis available to coalescent theory as applied to neutral evolution models and irreversible biologically motivated models typically simulated using time consuming Monte Carlo techniques.

Despite the advances presented here additional future work remains. The scaling exponent for the time to the most recent common ancestor in the Eden model appears to follow a generally accepted relationship with its dynamic exponent, but a comprehensive computational study of the tree structures in a more biologically realistic tissue models is clearly desirable. More importantly, however, many of the results presented here can be confirmed experimentally. In particular the determination of a dynamic exponent for an actual growing bacterial population combined with lineage tracing experiments might enable the reproduction of Figure 5 and confirm the computational results presented. Of particular appeal is the potential to estimate the number of stem-cells required to generate the cells in a tissue of given size over a certain limited time-frame. We hope that this study motivates experimental analyses that allow us to gauge the required size of the stem cell pool, as this (1) would be the most stringent test for our theoretical analysis, and (2) could have profound implications for developmental biology as well as regenerative medicine.

Already, however, our analytical framework provides a useful complementary framework for the analysis of lineage tracing studies. The results found in the Eden growth models suggest that many conclusions otherwise discovered through multiple costly ABM Monte Carlo simulations (such as “superstars” [16]) are readily explained by, and indeed expected from, coalescent theory. Finally, there is, we believe an intrinsic appeal of applying evolutionary concepts to developmental problems. Evolution does, of course, provide a framework against which we view development, but here it can also provide powerful computational tools for the analysis of tissue dynamics during growth as well as homeostasis.

## Acknowledgments

The authors gratefull acknowledge financial support from the *Biotechnological and Biological Research Council* through grant BB/K003909/1.

## Footnotes

1 These two terms have semantically diverged since the 1970s. A *complex system* is one where the interactions at lower levels lead to “emergent” behaviour at higher levels (e.g. behaviour that was not explicitly included in the rules governing behaviour at lower levels.)

